# Glucose oxidation and nutrients availability drive neural crest development

**DOI:** 10.1101/2022.09.05.506657

**Authors:** Nioosha Nekooie-Marnany, Redouane Fodil, Sophie Féréol, Marine Depp, Roberto Motterlini, Roberta Foresti, Jean-Loup Duband, Sylvie Dufour

**Affiliations:** Institut Mondor de Recherches Biomédicales, INSERM U955, Université Paris-Est Créteil, 94000 Créteil, France

**Author notes:** Institut de Biologie, Ecole Normale Supérieure, Paris, France (Marine Depp).

**Keywords:** neural crest, cellular metabolism, glycolysis, oxidative phosphorylation, nutrient input, bioenergetics, cell migration, fate decision

## Abstract

Bioenergetic metabolism is a key regulator of cellular function and signaling activity but the exact roles of nutrient utilization and energy production in embryonic development remain unknown. Here we investigated the metabolic pathways and deciphered the role of carbon metabolism required for the development of neural crest cells (NCC), a migratory stem cell population of the vertebrate embryo. We uncovered that glucose oxidation constitutes the prominent metabolic signature of trunk NCC and supports their delamination, migration, and proliferation. Additionally, we found that glycolysis, mitochondrial respiration and the pentose phosphate pathway are all mobilized downstream of glucose uptake. These metabolic pathways do not support specific cellular processes but cooperate and are integrated to accomplish epithelium-to-mesenchyme transition, adhesion, locomotion and proliferation. Moreover, using different nutrient supplies (glucose vs. pyruvate) we show that glucose is crucial to modulate NCC migration and adaptation to environmental stiffness, control NCC stemness and drive their fate decisions through regulation of specific gene expression. Our data establish that NCC development is instructed by metabolic cues that mobilize defined metabolic pathways cooperating together in response to nutrient availability.

**SUMMARY STATEMENT:** Here we show that neural crest cell migration and fate decisions rely primarily on glucose oxidation for energy production and mobilize multiple cooperating metabolic pathways for their biosynthetic needs and execution of gene programs.

## INTRODUCTION

During embryonic development, cell type diversity is generated through migration of progenitor cells over considerable distances and during long time periods. Neural crest cells (NCC) of the vertebrate embryo are one remarkable example of such migratory populations (Bronner, 2018; Trainor, 2013). These cells appear early all along the embryonic rostrocaudal axis in the dorsal part of the neural tube (NT). Through epithelium-to-mesenchyme transition (EMT), NCC delaminate and disperse away from the NT to distal sites where they differentiate. Owing to their partition into different populations (cranial, cardiac, vagal, trunk, and sacral) along the neural axis, NCC give rise to numerous cell types: skeletal and supportive tissues, endocrine cells, cardiac septum, enteric ganglia as well as peripheral neurons, glia, and melanocytes. From delamination to differentiation, NCC are confronted with different environments and are subjected to diverse cues impacting on gene networks and on their survival, proliferation, shape, adhesion, migration, and fate (Barriga and Mayor, 2019; Chevalier et al., 2016; Duband et al., 2015; Martik and Bronner, 2017; Soldatov et al., 2019).

Cellular metabolism, long regarded as a neutral factor during development, has been proposed recently to have an instructing role in cell fate (Miyazawa and Aulehla, 2018; Pavlova and Thompson, 2016; Shyh-Chang et al., 2013; Zhang et al., 2018), but the exact roles of nutrient utilization and energy production in embryonic development still remain to be investigated. The primary function of cellular metabolism is to provide cells with bioenergetic and biosynthetic supplies from nutrients, e.g. glucose, pyruvate, amino acids, and fatty acids (Zhu and Thompson, 2019). Glucose constitutes the major source for ATP production through two pathways occurring in distinct cellular compartments: glycolysis in the cytoplasm and oxidative phosphorylation (OXPHOS) in mitochondria. Glucose is also an important source of carbon to produce amino acids, lipids, and nucleotides necessary for biomass production and generated from metabolic pathways branching from glycolysis. Among them, the pentose phosphate pathway (PPP) constitutes a major source of nucleotides and NADPH required during anabolism (Patra and Hay, 2014).

Bioenergetic and biosynthetic metabolisms differ among cells and depend on gene regulatory programs driving expression and activity of metabolic enzymes (Lempradl et al., 2015). For example, in cell cycle-arrested differentiated cells, metabolism is usually catabolic with high production of ATP, as a result of pyruvate entry into mitochondria and OXPHOS. In contrast, in proliferating, undifferentiated cells as well as in cancer cells, metabolic activity is preferably oriented toward anabolism and ATP production relies mostly on aerobic glycolysis fed by intense glucose uptake, with high extracellular lactate production, a process known as Warburg effect (Schell et al., 2017; Vander Heiden et al., 2009). On the other hand, depending on the nature and amount of available nutrients, the metabolic activity of cells varies in time and space to adapt to environmental changes. For example, the mouse embryo relies essentially on OXPHOS before implantation, while after implantation aerobic glycolysis predominates until OXPHOS takes over again when blood flow begins (Johnson et al., 2003; Miyazawa and Aulehla, 2018; Perestrelo et al., 2018; Shyh-Chang et al., 2013; Zhang et al., 2018). A similar shift from glycolysis to OXPHOS takes place in adult stem cells leaving quiescence to differentiate and in induced pluripotent stem cells exiting pluripotency (Gu et al., 2016; Ito and Suda, 2014; Perestrelo et al., 2018; Shyh-Chang et al., 2013).

Recently, a study on cranial NCC delamination reported a metabolic shift toward aerobic glycolysis at initiation of their migration (Bhattacharya et al., 2020), suggesting that NCC can modulate their metabolic activity to execute different developmental steps. However, cranial NCC delamination and migration differ strikingly from those of the other NCC populations in their dynamics, gene regulatory network deployed, and environment (Li et al., 2019; Scully et al., 2016; Simões-Costa and Bronner, 2015; Simões-Costa et al., 2012; Soldatov et al., 2019; Théveneau et al., 2007). Therefore, a global view of the NCC metabolic landscape is still lacking, and it remains to be explored how nutrients support different NCC cellular activities throughout development, which metabolic pathways are recruited for their bioenergetic and biosynthetic supplies, and to which extend NCC metabolic activity can drive their behavior and fate.

Because investigating NCC metabolic features and their impact on NCC developmental program cannot be performed directly in intact embryos, we developed an alternative strategy based on *ex vivo* culture in defined conditions recapitulating the delamination, migration, and differentiation steps of NCC development, as well as their main features (timing and kinetics of cellular events, rate of proliferation, cellular interactions) (Duband et al., 2020). We characterized the metabolic traits of avian trunk NCC, and we modified the activity of some of the key metabolic enzymes or the nutrient supplies to decipher the role of carbon metabolism in NCC dispersion and differentiation. Our results indicate that, NCC have a metabolic signature characteristic of glucose oxidation and their development requires massive mobilization of the glycolytic, OXPHOS and PPP pathways cooperating together to meet their strong bioenergetic and biosynthetic needs and to adapt to a fast-changing environment. In addition, our data reveal that nutrient inputs and metabolic pathways can drive fate decisions in NCC through regulation of gene programs.

## RESULTS

We established avian trunk NCC in primary cultures from neural tube (NT) explants in a defined medium suitable to support their developmental steps, from delamination and dispersion to differentiation (Fig. S1), at rates and timings comparable to those achieved in classical media (Dupin et al., 2018; Rovasio et al., 1983; Santiago and Erickson, 2002). This medium, referred as to control medium, consisted of DMEM containing glucose, pyruvate, and glutamine at 5, 1, and 2 mM, respectively, as bioenergetic and biosynthetic nutrients and supplemented with only 1% serum to minimize the contribution of exogenous growth factors and fatty acids.

### Multiple Metabolic Pathways are Mobilized during Trunk NCC Dispersion

We first investigated whether the spread of NCC outgrowth relies on specific metabolic pathways (Fig. 1A-D). Inhibitors of glycolysis, OXPHOS, and PPP (Fig. 1A) were applied at onset of culture (except in video microscopy analyses, where they were applied just before recording), and the behavior of the NCC population over 24 h was compared to that of explants in control medium, focusing on outgrowth aspect and area at 5 and 24 h (Fig. 1B,C), migration front progression (Fig. 1D) and morphology of cells (Fig. S2A). In the presence of 2-deoxyglucose (2-DG), a potent inhibitor of upper glycolysis, NCC initially segregated normally from the NT and their progression was close to that of controls, but cells became less spread and less cohesive, with a spindle shape. Then, expansion of the population almost completely stalled and NCC started to round up. At 24 h cells displayed poor survival and were sparse occupying a limited area around the NT, compared with the large outgrowth with numerous well-spread and dense cells seen in controls. Oligomycin, an inhibitor of mitochondrial ATP synthase, marginally affected initial NCC progression, but cells were densely packed, causing a reduction of the outgrowth area. Then, NCC expansion decreased with a progressive reduction in cell spreading, resulting in a reduced outgrowth of sparse, round cells at 24 h. Rotenone combined with antimycin-A (Rot-AA), blockers of complex I and III of the mitochondrial electron transport chain, had a similar effect except that reduction in dispersion was immediate and massive. 6-aminonicotinamide (6-AN), a blocker of PPP, showed no apparent effect on the initial dispersion of NCC, but the progression of cells slowed down after 5-8 h, and a strongly reduced outgrowth with many round cells was observed at 24 h. Na-iodoacetate, an inhibitor of lower glycolysis (Fig. S3A), reduced significantly NCC dispersion after 24 h but less dramatically than 2-DG (Fig. S3B). Unlike 2-DG, Na-iodoacetate did not alter cell survival but produced sparse outgrowths containing round to elongated NCC.

**Fig. 1.**
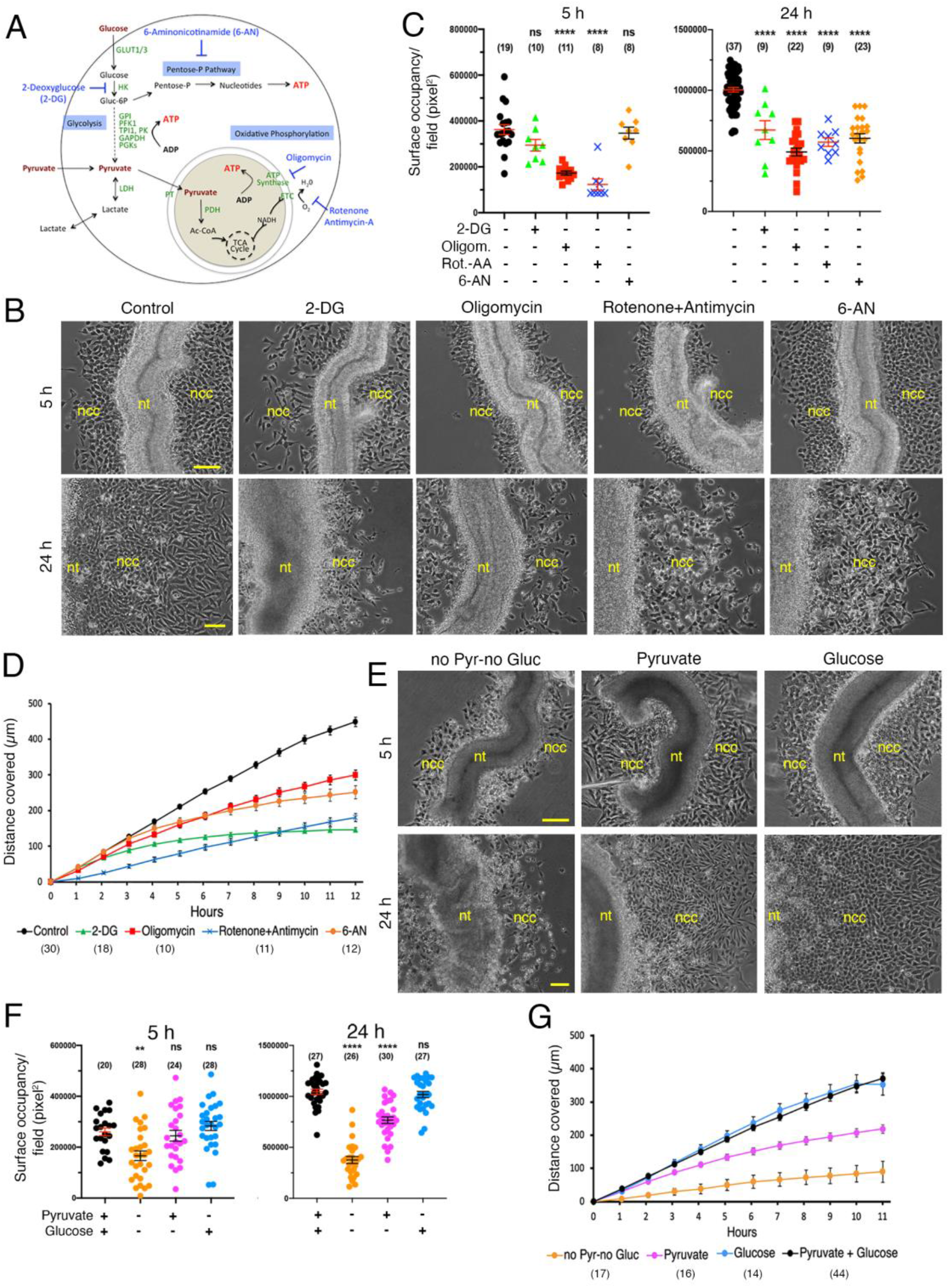
Multiple metabolic pathways are mobilized during trunk NCC dispersion. (A) Schematic representation of the main pathways of glucose and pyruvate metabolism indicating the specific targets of the respective inhibitors of glycolysis (2-DG), ATP synthase (oligomycin), complex I and III of the electron transport chain (Rot-AA), and PPP (6-AN). (B-G) Effect of metabolic inhibitors (B-D) and of nutrients on NCC dispersion (E-G): (B, E) Overall aspect of NT explants; (C, F) NCC outgrowth area; (D, G) Progression of the migratory front of the NCC population over time. Bars in B and E = 100 μm. Error bars = S.E.M.

Thus, all inhibitors of glucose metabolism and OXPHOS affected NCC dispersion even if with specific kinetic and morphological effects, suggesting that all metabolic pathways downstream of glucose uptake and utilization are mobilized in a cooperative manner in NCC.

### Nutrient Availability Affects Trunk NCC Dispersion

To investigate whether NCC development is influenced by nutrient availability, we cultured NT explants in conditions in which, beside glutamine, only pyruvate, glucose, or neither were provided (Figs. 1E-G, S2B). In the absence of pyruvate and glucose, NCC initiated migration from the NT but failed to follow their normal progression and dispersion over time, with considerable cell death after 24 h. In the presence of pyruvate, NCC dispersion occurred almost normally at culture onset but stopped after 5-8 h, with cells exhibiting elongated spindle shaped morphologies, thus mimicking strikingly the behavior and morphology of NCC treated with 2-DG, except that they did not undergo cell death in the long term. In the presence of glucose, progression of NCC outgrowths and cell morphologies were virtually indistinguishable from those observed in controls. Oligomycin exerted a much more immediate and stronger effect on NCC cultured in pyruvate medium than in glucose, with the abrogation of cell dispersion (Fig. S2C). Conversely, 6-AN greatly affected NCC dispersion in glucose medium but had little effect in pyruvate even after 24 h (Fig. S2C). UK-5099, a potent inhibitor of the mitochondrial pyruvate carrier (Fig. S3A) (Schell et al., 2017) reduced NCC dispersion and affected their spreading, severely in the presence of pyruvate and more moderately with glucose (Fig. S3C-F). We also observed that nutrient concentration affects NCC dispersion. Pyruvate at 1 mM and glucose at 5 mM present in the control medium were the most suitable conditions for NCC dispersion while either lower or higher concentrations of these nutrients failed to produce large NCC outgrowths (Fig. S4A). In addition, low concentrations of pyruvate did not support NCC spreading while at high concentrations it induced extensive NCC flattening (Fig. S4B). Glucose, in contrast, favored NCC spreading at all concentrations used (Fig. S4B).

All together, these results suggest that NCC dispersion relies strongly on nutrient inputs and downstream metabolic pathways. Specifically, glucose oxidation through glycolysis coupled to OXPHOS and PPP plays a pivotal role in dispersion. Pyruvate alone, which feeds only OXPHOS, is in contrast less favorable in the long-term.

### Metabolic Profiling of NCC Reveals a Typical OXPHOS Signature during Migration

We then assessed cellular bioenergetics in NCC and measured mitochondrial respiration and glycolysis using a Seahorse XF analyzer. Analyses of the bioenergetic profiles of more than 300 individual NT explants combined with MitoStress assays revealed that a great majority of the explants displayed an OXPHOS signature characterized by high oxygen consumption rate (OCR) and low extracellular acidification rate (ECAR) at 5 h (Fig. 2A), without notable changes at 24 h (Fig. 2B), suggesting that their metabolic activity was constant throughout migration. 2-DG or oligomycin caused a sharp drop in OCR with no significant alteration in ECAR (Fig. 2C-E), and they strongly reduced ATP production (Fig. 2F), but had no effect on the extracellular lactate level (Fig. 2G). In contrast, 6-AN had no effect on OCR and ATP levels (Fig. 2C-F). Similar to the control condition, NT explants exposed to glucose or pyruvate only were sensitive to MitoStress (Fig. 2H). Interestingly, lower values of OCR and ECAR and ATP production were observed in pyruvate condition compared to glucose (Fig. 2H-K). 2-DG and oligomycin decreased OCR levels (Fig. 2I, J) and sharply reduced ATP production in both media (Fig. 2K), but they did not affect lactate production (Fig. 2L). Thus, a better yield in energy production is obtained for glycolysis coupled to OXPHOS than for OXPHOS fueled by pyruvate alone. Importantly, our results demonstrate that, in explants cultured with control medium, most of the ATP production relies primarily on glucose oxidation through OXPHOS, and suggest that OXPHOS inhibition does not cause metabolic rewiring to aerobic glycolysis with increased lactate production.

**Fig. 2.**
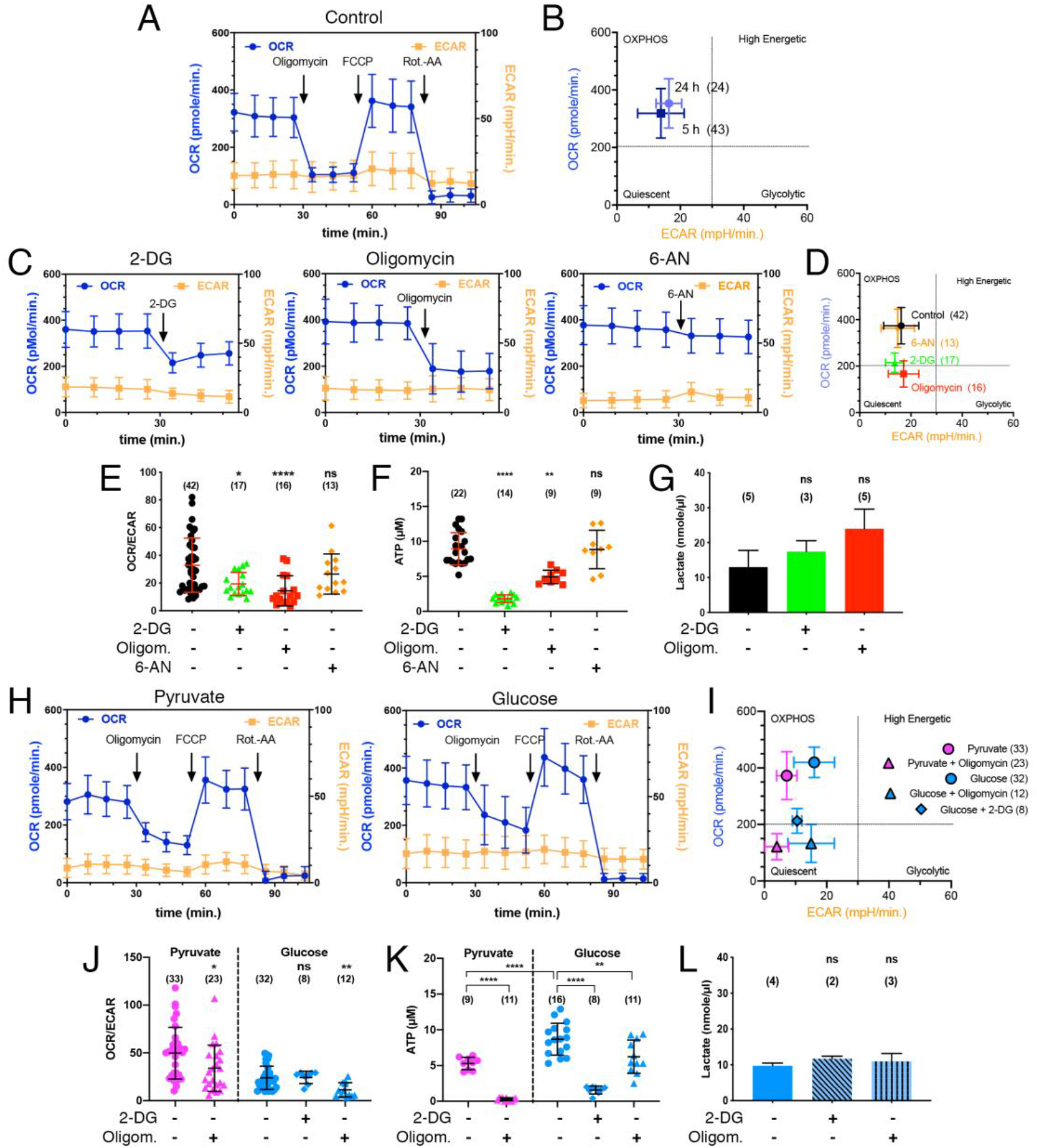
Metabolic profiling of NCC reveals a typical OXPHOS signature during migration. Quantitation analyses of O_2_ consumption rate (OCR) and extracellular acidification rate (ECAR) using a Seahorse analyzer, and ATP and lactate productions in individual NT explants cultured for 5 h (and 24 h in B) in control medium after treatment with metabolic inhibitors, and in different nutrients: (A, H) MitoStress assays; (B, D, I) Energetic maps deduced from OCR and ECAR measurements; (C) OCR and ECAR profiles over time in explants cultured in control medium in the presence of metabolic inhibitors; (E, J) Values of OCR/ECAR ratio; (F, K) Intracellular ATP measurement; (G, L) Extracellular lactate measurement. Error bars = S.D.

Finally, we analyzed the topological features of mitochondria network in NCC (Fig. S5). While in NCC cultured in glucose or in control medium, mitochondria had a large globular structure and were organized as rings homogenously distributed throughout the cell body, in pyruvate medium, they were smaller, elongated, and often organized as chains extending into the cell processes (Fig. S5A). As expected, mitochondria were small and poorly organized in the absence of nutrients (Fig. S5A), as observed in the presence of oligomycin or UK-5099 (Fig. S5B), further illustrating the critical role of mitochondria in NCC.

### Expressions of Glucose Transporter and Glycolysis Enzymes are Regulated at the Time of Migration

To probe the link between nutrient supply and metabolic genes expression, we analyzed the expression of key players of glycolysis in NCC. The glucose transporter Glut-1 was expressed equally on the surface of NCC in pyruvate and glucose conditions (Fig. 3A). *Glut-1* mRNA as well as those for phosphofructokinase *PFK*, a major rate-limiting enzyme of upper glycolysis (Feng et al., 2020), were clearly expressed in the NT but barely detected in NCC. In addition, they were both up regulated in the presence of glucose compared to pyruvate (Fig. 3B,C), indicating that their expression relied on nutrient supply. In contrast, phosphoglycerate kinase-1 (*PGK-1*), an enzyme of lower glycolysis, was expressed in both the NT and migrating NCC and did not show much differential expression with nutrients (Fig. 3B). In embryos, *Glut-1* and *PFK* genes were poorly expressed in the caudal trunk region except in the NT in which they were strongly but transiently up regulated at the time of NCC delamination (Fig. 3D). *PGK-1* was expressed in all tissues, but was increased in the NT during delamination.

**Fig. 3.**
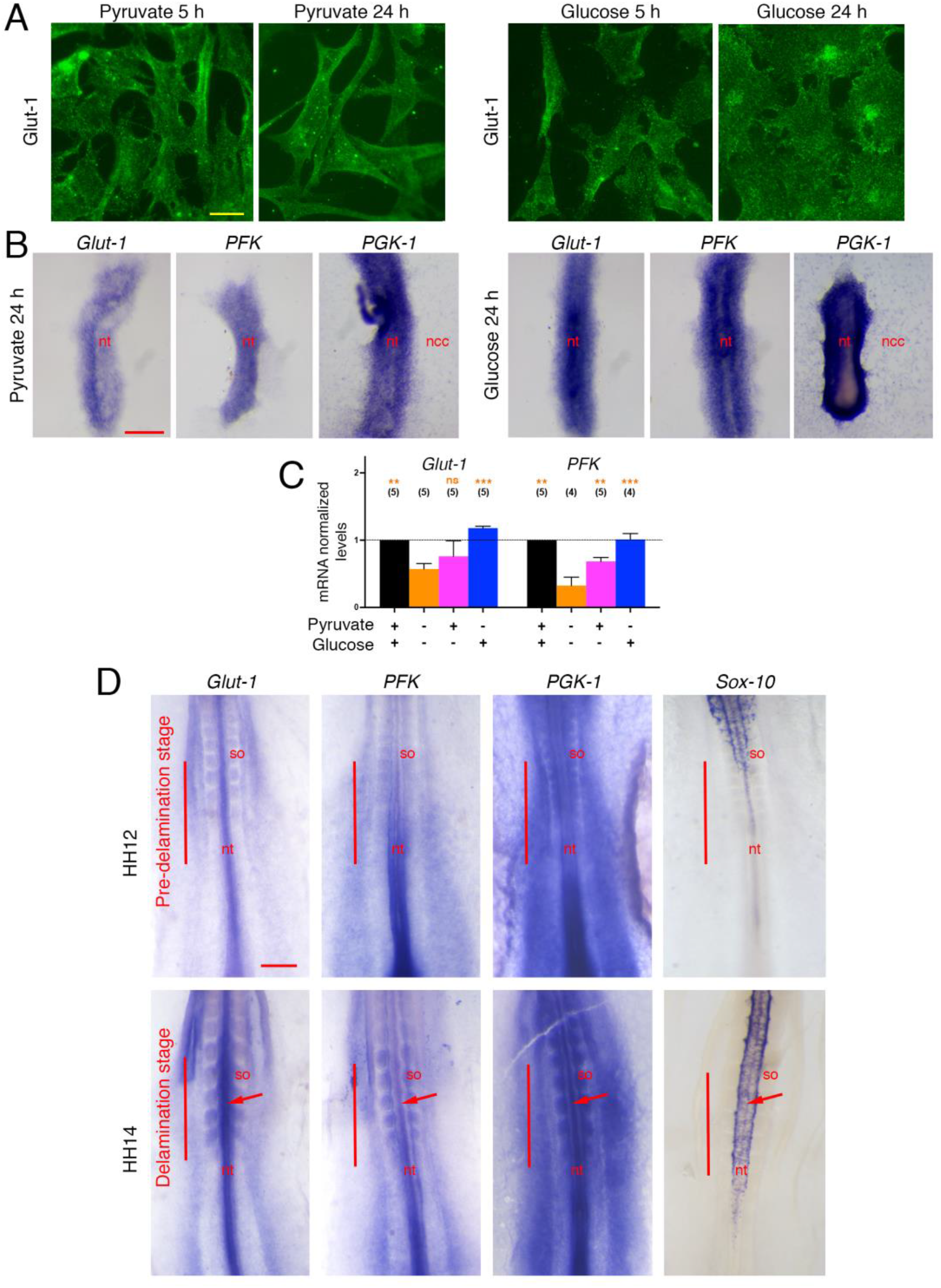
Expressions of glucose transporter and glycolytic enzymes are regulated at the time of NCC migration. (A-C) Immunostaining (A), in situ hybridization (B), and qPCR (C) analyses of the expression of the glucose transporter *Glut-1* and of glycolytic enzymes *PFK* and *PGK-1* in NCC and NT explants. (D) Whole mount in situ hybridization for *Glut-1, PFK*, and *PGK-1* and for the migratory NCC marker *Sox-10* in avian embryos at stage HH12 and 14 (i.e. 17 and 22 somites, respectively), showing the caudal part of the embryo at the level of the last formed somites (so) and of the unsegmented mesoderm. The bars indicate the axial levels at which NCC are in the pre-delamination stage at HH12 and in the delamination stage at HH14. Arrows point at the strong staining for *Glut-1, PFK*, and *PGK-1* in the NT at levels of delaminating NCC, contrasting with the fainter NT staining in regions of pre-delaminating NCC. Bars in A = 10 μm, in B = 100 μm, and in D = 200 μm. In C, comparisons were done respective to the no nutrient condition. Error bars = S.E.M.

These data indicate that while lower glycolysis is constantly active in migrating NCC, upper glycolysis is subjected to spatial and temporal regulation at onset of migration, therefore raising the intriguing possibilities that, in migrating NCC, lower glycolysis may be fed by exogenous metabolic intermediates possibly provided by the NT; and localized activation of glycolytic enzymes in tissues may be due to differential distribution of nutrients and metabolites in the embryo.

### Glucose Metabolism Regulates *Snail-2* Expression during Delamination

NCC dispersion results from multiple cellular processes acting in synergy and in concerted manner: EMT during delamination, changes of substratum and cell adhesion, activation of the locomotory machinery, and proliferation. Since higher expression of metabolic enzymes was observed upon NC emergence, we studied the role of metabolic pathways from the delamination to the migration step and developed an assay suitable for discriminating delaminated from migrating NCC on 5h NT explants (Fig. 4A). Interestingly, the number of delaminated cells was halved by addition of 2-DG, oligomycin, or Rot-AA, as well as in pyruvate medium, while in the presence of 6-AN, it was similar to control (Fig. 4B). Furthermore, Snail-2, a major EMT player (Duband, 2010; Nieto, 2002) was high in delaminated NCC but declined gradually as cells dispersed away from the NT in control and in glucose or 6-AN conditions, while in 2-DG or pyruvate, it was strongly diminished in both NCC compartments (Fig. 4C). In contrast, consistent with an unchanged delamination rate in response to 6-AN, NCC displayed similar Snail-2 staining under PPP inhibition (Fig. 4C). Intriguingly, with OXPHOS inhibitors and particularly with Rot-AA, Snail-2 completely disappeared in delaminated NCC but remained high in migrating cells. Expression of *Snail-2* messengers in NT explants (Fig. 4D,E) was almost totally abrogated by 2-DG and strongly diminished by oligomycin, and was low in pyruvate and no nutrient conditions, consistent with the reduction of Snail-2 protein. Conversely, *Snail-2* expression was not much altered by Rot-AA, probably as a result of its increase in the migrating pool.

**Fig. 4.**
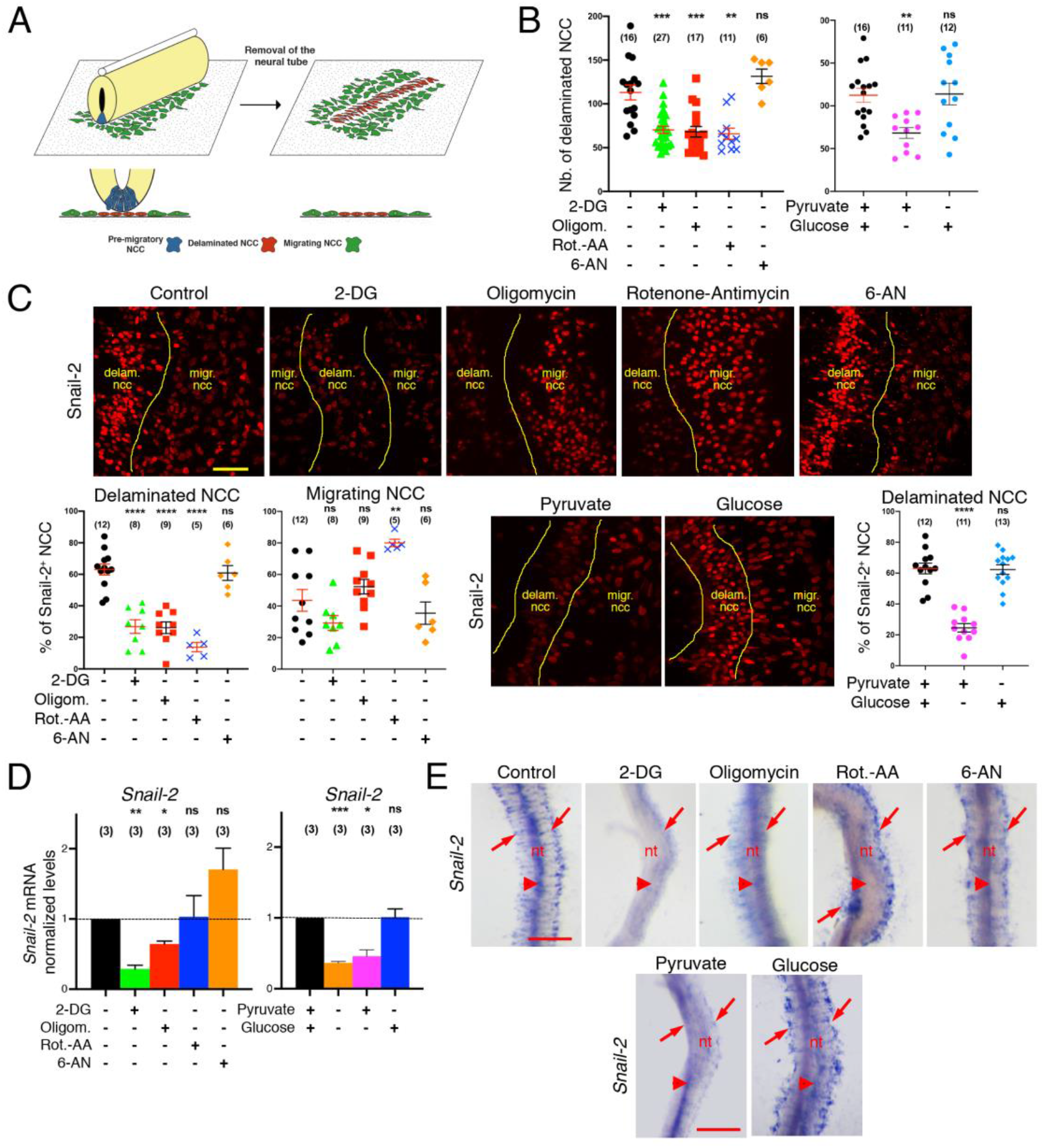
Glucose metabolism regulates *Snail-2* expression during NCC delamination. (A) Schematic representation of the delamination assay. (B, C) Effect of metabolic inhibitors and of nutrient supplies on the number of delaminated cells per explant (B) and expression of the Snail-2 transcription factor (C) in delaminated vs. migrating NCC. (D, E) qPCR (D) and in situ hybridization (E) analyses of *Snail-2* expression in NT explants. In control medium, in the presence of 6-AN or with glucose, *Snail-2* messages are distributed in premigratory NCC in the dorsal NT (E, arrowheads) as well as in the delaminated NCC visible in either sides of the NT (E, arrows). In the presence of 2-DG, oligomycin, and in pyruvate medium, *Snail-2* is much decreased in both premigratory and migratory NCC while in the presence of Rot-AA, it drops only in premigratory cells. Bars in C = 50 μm and E = 100 μm. Error bars = S.E.M.

All together, these data indicate that glucose metabolism is required for NCC delamination by regulating the coordinated expressions of EMT-related genes and that OXPHOS under pyruvate condition is necessary but not sufficient for this process.

### Glucose Metabolism is Crucial for Locomotion

Avian trunk NCC display a locomotory behavior different from the ones described for cranial or enteric NCC populations (Genuth et al., 2018; Kulesa and Fraser, 1998; Li et al., 2019; Rovasio et al., 1983). Movement of individual cells involves substrate adhesion and intense protrusive activity, while outward directionality is driven by transient cell adhesion and mechanical forces resulting from high cell density. To better understand the contribution of glucose metabolism to NCC locomotion, we analyzed by video microscopy the effect of metabolic inhibitors applied after initiation of NCC migration to avoid any bias due to the delamination process (Figs. 5A,B, S3G,H, S6A,B; see also Figs. 1, S2). In controls, NCC generally exhibited persistent and directional migration oriented perpendicular to the NT leading to relatively linear trajectories (Fig. S6A). Both the velocity and persistence of cells were constant over 18 h, due to maintenance of high cell density (Fig. 5A,B). With 2-DG, cells trajectories were at first relatively linear but became more random with time. The most striking effect of 2-DG, however, was a rapid and continuous decrease of velocity over time and a progressive rounding up, resulting in migration arrest. Unlike 2-DG effect, cell trajectories were generally not much affected by oligomycin, while velocity was decreased during the first 6 h. Subsequently cell migration became extremely random with decreased persistence and higher velocity linked to gradual rounding up of the cells and lack of cell-cell contacts. Interestingly, Rot-AA showed a much stronger effect than oligomycin. Trajectories were very short because velocity was almost knocked-out after 2 h and remained weak during 8 h. However, as with oligomycin, velocity increased with time and persistence decreased coincident with cell rounding up. In UK-5099, cells trajectories were more random, associated with low persistence (Fig. S3G,H). Lastly, 6-AN affected velocity strongly and gradually during the first 8 h before stabilizing, and cell trajectories became less linear (Figs. 5A,B, S6A). NCC locomotion was also impacted by nutrient supply (Figs. 5C, S4C,D, S6B). In absence of nutrients, velocity was consistently low with good persistence, resulting in very short migration tracks. In glucose condition, NCC displayed long trajectories, with high and sustained velocity and persistence over time as seen in controls. In contrast, in pyruvate medium, NCC trajectories exhibited a profile similar to those seen with 2-DG, with a gradually decreasing velocity and reduced persistence (Fig. 5C, S6B). Velocity peaked for pyruvate at 1 and 2 mM and plateaued for glucose concentrations ≥ 5 mM (Fig. S4C), in agreement with the front progression curves (Fig. 1G). Persistence, in contrast, was greater for pyruvate concentrations ≥ 2 mM and constant for glucose concentrations > 1 mM, possibly reflecting the differences observed in cell spreading (Fig. S4B).

**Fig. 5.**
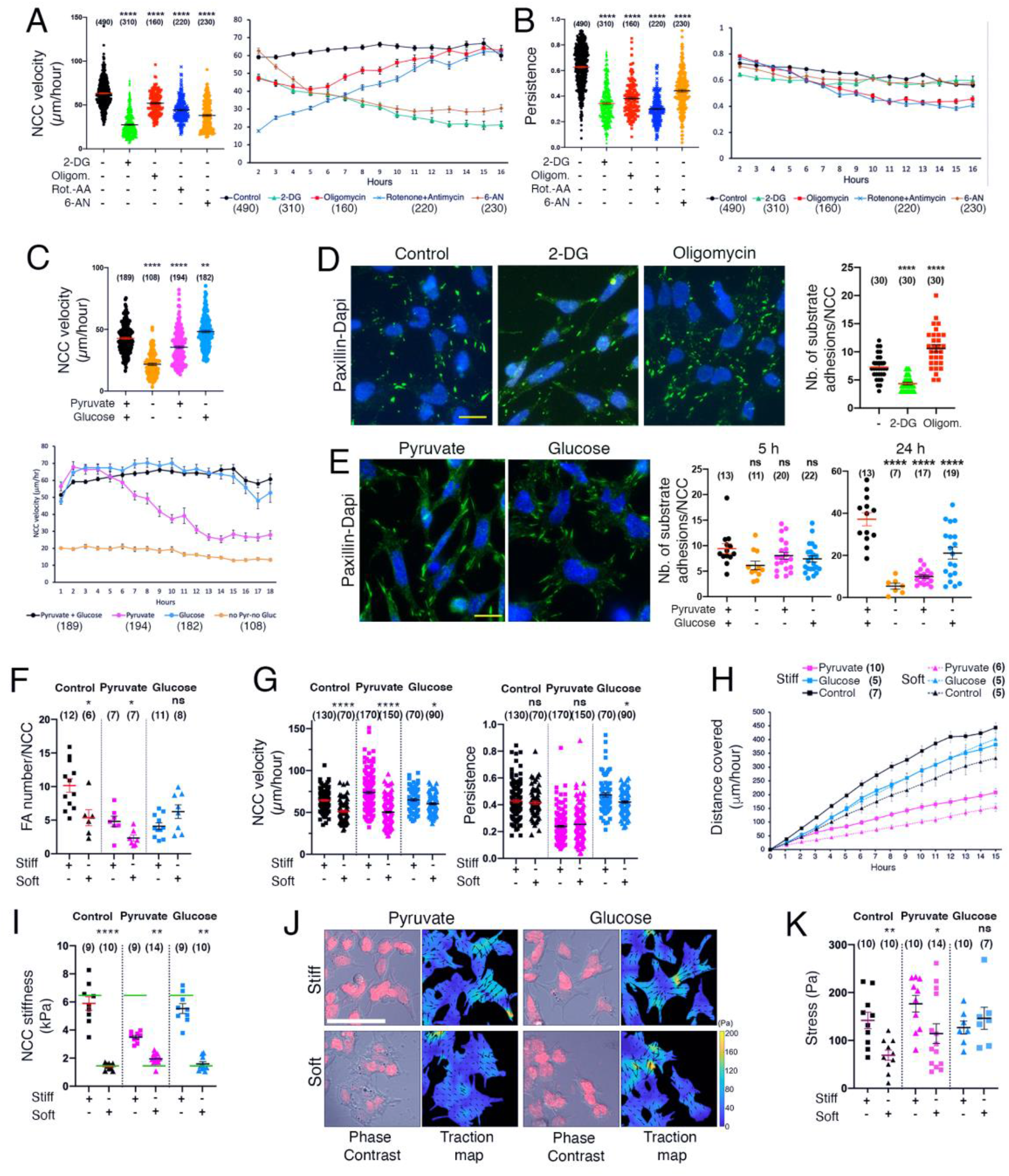
Glucose metabolism is crucial for NCC locomotion and regulates NCC adhesion and mechanical properties and response to External Stiffness. (A-E) Effect of metabolic inhibitors and of nutrient supplies on NCC locomotion. (A, C) Velocity of individual NCC throughout the duration of the video microscopy recording and variation of NCC velocity over time in culture. (B) Persistence of migration of individual NCC throughout the duration of the video microscopy recording and variation of the persistence over time in culture. (D) Effect of metabolic inhibitors and (E) of nutrients on NCC substrate adhesion. Visualization of paxillin (green) and Dapi (blue) expression and quantification of the number of substrate adhesions in migrating NCC. (F-H) Effect of nutrient supplies on NCC adhesion and migration on stiff and soft gels in 5-h cultures. (F) Quantification of substrate/focal adhesions (FA) number per NCC. (G-H) Cell locomotion; velocity (G, left panel) and persistence of migration (G, right panel) of individual NCC throughout the duration of the video microscopy recording. (H) Progression of the migratory front of the NCC population over time. (I-K) Effect of nutrient supplies on NCC mechanical properties. (I) NCC stiffness on stiff and soft gels. The green bars indicate the stiffness of the gels. (J) Phase contrast and traction map images. (K) Traction activity of NCC expressed as stress per NCC. Bars in D, E = 20 μm, and in J =50 μm. Error bars = S.E.M.

These results therefore illustrate the critical contribution of glucose metabolism to NCC locomotion and also reveal that this process relies on other metabolic processes, notably PPP as well as non-energetic mitochondrial activities involving TCA metabolites and byproducts.

### Glucose Metabolism Coordinates Substrate and Cell Adhesions

To gain insight into the impact of metabolism on cell adhesions, two pivotal aspects of NCC delamination and migration, we examined the cellular localization of paxillin and β-catenin, two cytoplasmic partners of integrins- and cadherins-adhesion complexes, respectively (Figs. 5D,E, S6C,D). Compared to controls, at 5 h, the number of substrate adhesions per NCC was significantly reduced with 2-DG and increased with oligomycin, consistent with changes in cell spreading (Fig. 5D). In addition, oligomycin increased substantially cell-cell adhesions during the first 5 h (Fig. S6C). The number of substrate and cell-cell adhesions depended on the nutrient supplies (Fig. 5E, S6D). In control and glucose medium, NCC exhibited large substrate adhesions with a four- and two-fold increase in their number at 24 h, respectively (Fig. 5E). In contrast, in pyruvate medium, NCC displayed more elongated substrate adhesions, which did not increase in number with time. In no nutrient condition, NCC developed substrate adhesions at 5 h but their number decreased with time. Quantification of cell-cell adhesions revealed that in control as well as in glucose medium, NCC are always engaged in transient contacts with 1-3 neighbors while in pyruvate, they remain as individuals (Fig. S6D).

These data indicate that glucose metabolism regulates the balance between substrate and cell-cell adhesion in NCC, thereby favoring optimal conditions for delamination and migration.

### NCC Mechanics and Response to External Stiffness are Modulated by Nutrient Inputs

NCC are able to sense the extracellular matrix rigidity in their environment, which in turn modulates their adhesion and migration capacity (Chevalier et al., 2016; Shellard and Mayor, 2021). To determine whether mechanotransduction activities in NCC are under metabolic control, we investigated the impact of nutrients on their adhesive and migratory response to fibronectin-coated polyacrylamide gels with different stiffness in the range of tissue rigidity at early embryonic stages, from soft (1.2 kPa) to stiff (6.3 kPa) (Chevalier et al., 2016; Discher et al., 2005). We first characterized and quantified NCC substrate adhesions (Fig. 5F, S6E), velocity and persistence (Fig. 5G), and migration front progression (Fig. 5H). As on glass (Fig. 5D), NCC on stiff and soft gels developed substrate adhesions irrespective of the nutrient supplied (Fig. S6E), Moreover, we found important discrepancies in NCC response to gel rigidity with the nutrients (Fig. 5F-H). In glucose medium, the number of substrate adhesions in individual NCC and the progression of the population were not significantly different between stiff and soft gels and individual cell velocity and persistence were only slightly lower on soft gels. In contrast, in pyruvate and control medium NCC displayed less substrate adhesions on soft gels than on stiff gels, and their velocity and the progression of the population were significantly lower.

We then studied NCC stiffness and traction activity on the gel using atomic force and traction force microscopy (AFM, TFM) (Fig. 5I-K). Interestingly, measurement of cellular stiffness on gels revealed that in glucose and control medium NCC adapted their stiffness to that of the substrate in both soft and stiff gels, whereas in pyruvate they failed to increase their stiffness to the level of the gel when confronted with stiff gel (Fig. 5I). Comparison of NCC traction activity in soft and stiff gels expressed as force or stress/cell showed that it was constant in glucose whereas in pyruvate and control medium it was higher in stiff gels, though very heterogeneous (Fig. 5J, K)). Finally, by calculating the product of the average force and the average displacement per hour, we estimated the energy produced during NCC displacement on the soft and stiff gels under the different nutrient conditions. Energy produced was similar in all nutrients on stiff gels, while it was lower on soft gels in control and pyruvate medium compared to glucose in which it was almost unchanged (Table I).

**Table I.**
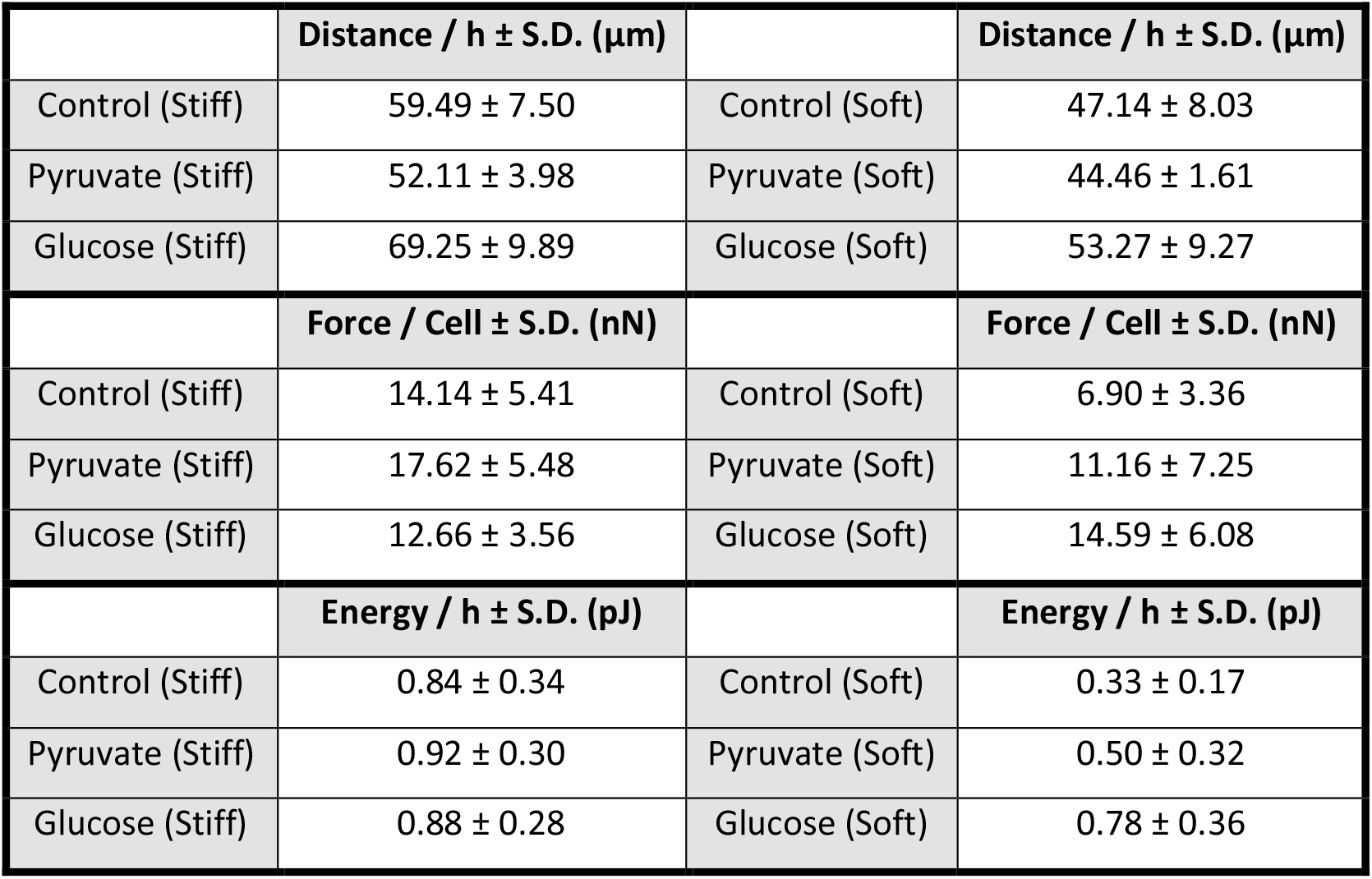
Energy produced by NCC on soft and stiff gels in various nutrients. Energy (expressed in pJ per h) was calculated as the product of the average force/cell in nN obtained by TFM and the average distance covered per hour in μm. P values for control, pyruvate and glucose between stiff and soft gels are 0.016, 0.063 and 0.636, respectively.

These data indicate that trunk NCC are mechano-responsive as described previously for cranial and enteric NCC (Chevalier et al., 2016; Shellard and Mayor, 2021), but that their response is determined by specific metabolic activities. In glucose medium, owing to their capacity to accommodate their stiffness and energy production to their environment, NCC are endowed with the ability to migrate in a great diversity of biophysical constraints. In pyruvate medium, NCC are less prone to modulate their stiffness and negatively respond to modifications in the rigidity of their environment by decreasing their adhesive and migratory response.

### Glucose Metabolism is Required for Sustained Proliferation

Contrary to many motile cells, NCC maintain active cell division throughout migration, and although this process by itself is not the driving force of migration, it is required for delamination (Burstyn-Cohen and Kalcheim, 2002) and contributes strongly to expansion of the population (Ridenour et al., 2014). We studied the impact of metabolic inhibitors on NCC proliferation. While, in control numerous proliferating (EdU^+^) NCC could be identified during their delamination along the NT apical side, only few proliferative cells were observed with glycolysis or OXPHOS inhibitors (Fig. S7A). These results indicate the contribution of glycolytic and mitochondrial intermediates in NCC proliferation during delamination. During migration, all inhibitors decreased both the proportion of EdU^+^ NCC and the EdU-staining intensity (Fig. S7B). Interestingly, unlike other compounds, the impact of 6-AN on proliferation decreased over time, possibly reflecting the contribution of other metabolic pathways to the adaptation to PPP inhibition. With no nutrient, proliferation was low at 5 h and was almost abolished after 24 h. Interestingly, consistent with the kinetics of their dispersion (see Fig. 1G), NCC proliferation observed in glucose was similar to that of controls whereas it was dramatically reduced after 24 h in pyruvate (Fig. S7C).

These data reveal that all metabolic pathways are implicated in a cooperative manner in the control of NCC proliferation.

### Pluripotency and Fate Decision are Dictated by Exogenous Nutrient Inputs

To investigate whether metabolic activity affects NCC stem cell capacity and influences their fate, we analyzed the expression patterns of the transcription factors Sox-10 and Foxd-3, two essential regulators of NCC pluripotency (Lukoseviciute et al., 2018; Schock and LaBonne, 2020; Simões-Costa et al., 2012). At 5 h in control, the vast majority of migrating NCC as well as some delaminating cells exhibited Sox-10^+^ nuclei, a pattern that mirrors that of Snail-2 (Fig. 6A, compare with Fig. 4C). 2-DG strongly diminished the number of Sox-10^+^ cells leaving a few remaining strongly-expressing cells while oligomycin and Rot-AA caused an overall reduction of Sox-10 staining intensity in all NCC (Fig. 6A). Likewise, *Sox-10* and *Foxd-3* mRNA levels dropped strongly with 2-DG and more moderately with oligomycin and Rot-AA (Fig. 6B). In addition, *Foxd-3* expression was selectively repressed by 2-DG, oligomycin or Rot-AA in pre-migratory NCC but less so in delaminating and migrating cells (Fig. 6C). Levels of *Sox-10* and *Foxd-3* messages and proteins were also much reduced in delaminating and migrating NCC cultured in pyruvate medium or with no nutrient compared with glucose (Fig. 6A-C). At 24 h (Fig. S8A,B), repression of *Sox-10* and *Foxd-3* expression was even more pronounced in the presence of metabolic inhibitors, including 6-AN, or in absence of glucose. These results indicate that sustained glucose metabolism is required for maintenance of high levels of pluripotency markers during NCC migration.

**Fig. 6.**
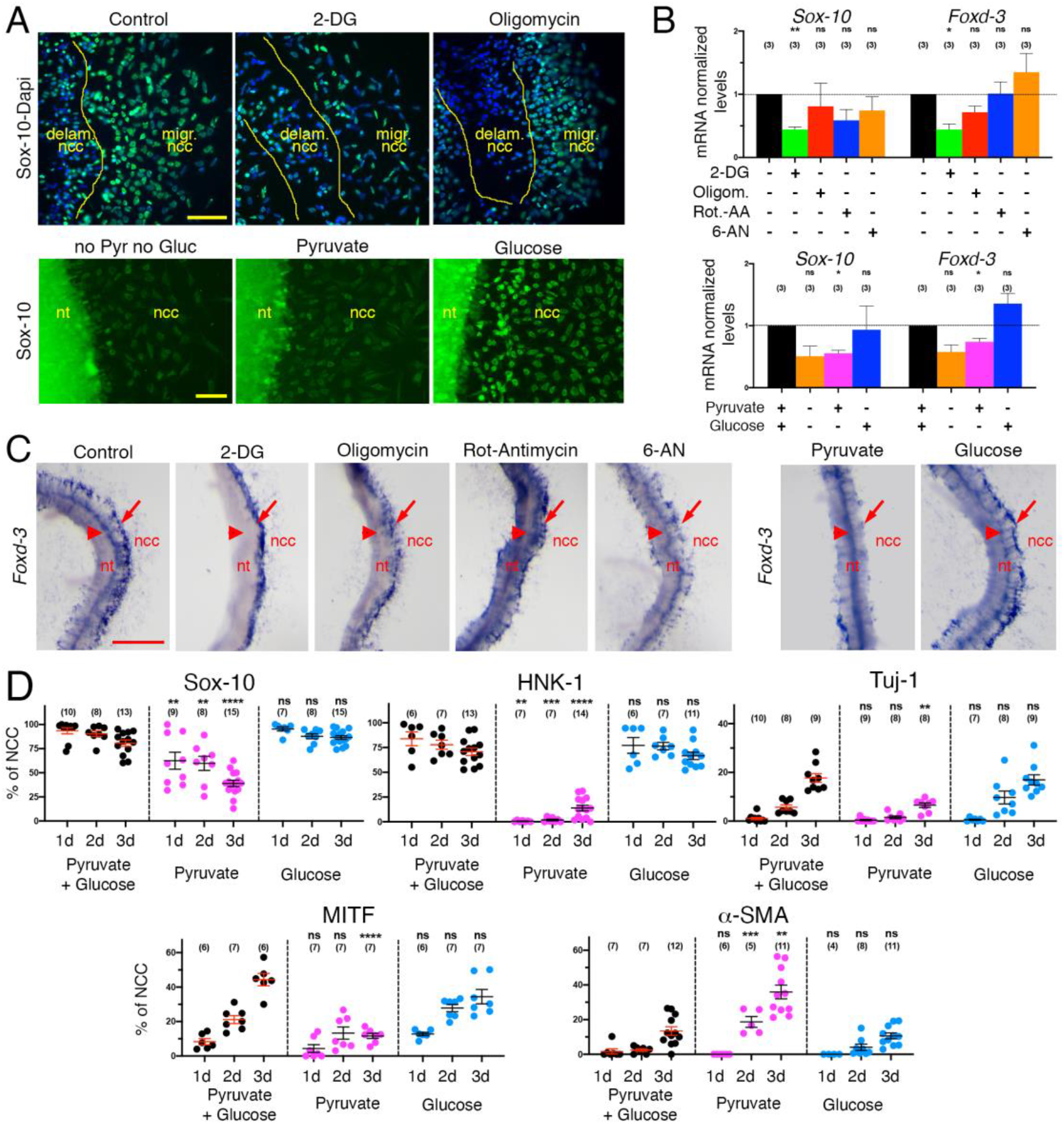
Glucose metabolism and nutrients supply regulate NCC pluripotency and fate decisions. Immunostaining for Sox-10 (A), qPCR (B), and in situ hybridization (C) analyses of the expression of *Sox-10* and *Foxd-3* transcripts in delaminating and migrating NCC after 5 h. In A, nuclei are visualized with Dapi. In control and glucose medium, *Foxd-3* messages are distributed in premigratory NCC in the dorsal NT (C, arrowheads) as well as in the delaminated cells visible in either sides of the NT (C, arrows) similar to that of *Snail-2*. In 2-DG and oligomycin, it is decreased essentially in premigratory cells whereas in pyruvate medium it is reduced only in delaminating cells. (D) Quantification of the proportion of Sox-10^+^, HNK-1^+^, Tuj-1^+^, MITF^+^, and α-SMA^+^ cells in NCC primary cultures over time under different nutrient conditions. Bar in A = 50 μm. Error bars = S.E.M. Bars in A = 50 μm, and C = 100 μm. Error bars = S.E.M.

We next searched for the appearance of differentiated cells in culture over time and analyzed the kinetics of expression of markers of NCC differentiation into the neural, melanocytic, and myofibroblastic/mesenchymal lineages. Because of the long-term deleterious effects of the metabolic inhibitors and lack of nutrient on NCC, we only examined cells cultured in the presence of pyruvate, glucose or both (Fig. 6D, S1C, S8C). In pyruvate medium, expansion of the NCC population stalled during the first day of culture and cell morphologies gradually shifted predominantly to a large, flattened fibroblastic shape after 3 days (Fig. S8C). In glucose medium in contrast, the NCC population steadily expanded, both in the cell number and in the area occupied, accompanied by the appearance of neurons and other cells presenting a great diversity of morphologies. In addition, the number of cells with a NCC shape did not decrease as dramatically as in the presence of pyruvate (Fig. S8C). In glucose medium, the proportion of NCC progenitors (Sox-10^+^ and HNK-1^+^) remained high over time (> 75%) while the number of neurons (Tuj-1^+^, Sox-10^-^ and HNK1^+^) and melanoblasts (MITF^+^, Sox-10^+^ and HNK1^-^) increased up to 20% (Fig. 6D). In pyruvate medium, the proportion of NCC progenitors (Sox-10^+^) dropped to less than 50% at the benefit of myofibroblasts (α-SMA ^+^ and Sox-10^-^), which peaked to about 40%. Of note, the number of melanoblasts did not increase much in pyruvate medium while the number of HNK1^+^ cells was constantly very low. A likely explanation for this persistent absence of HNK-1 staining is that the HNK-1 epitope is a glycoconjugate moiety and needs glucose for its biosynthesis (Tucker et al., 1984).

Our results indicate that NCC differentiation programs are dependent on nutrient inputs, with glucose favoring neural and melanoblast differentiation, while pyruvate promotes a limited range of phenotypes where myofibroblasts are predominant.

## DISCUSSION

Our study gives a general view of the metabolic impacts on NC development. We report that trunk NCC display a typical OXPHOS signature and that glucose oxidation through glycolysis coupled to OXPHOS plays a critical role in NCC dispersion. Blockers of glycolysis and OXPHOS affected every aspect of NCC development and caused alterations of mitochondria morphology associated with a sharp drop in mitochondrial respiration and ATP production. Additionally, our data establish that, though necessary for trunk NCC development, OXPHOS is not sufficient to support all the cellular events involved in this process as systematic inhibition of glycolysis compromised NCC dispersion, even in the presence of pyruvate to support OXPHOS. Likewise, a PPP inhibitor strongly affected NCC dispersion despite a minimal effect on mitochondrial respiration and ATP production. Finally, pyruvate alone failed to support complete dispersion, division, survival, and differentiation of NCC, while glucose could elicit the complete cellular response of NCC, i.e. delamination, adhesion, migration, proliferation, maintenance of stemness, and widespread differentiation. Glucose also potentiates NCC stiffness adaptation to that of their microenvironment and supports optimal dispersion, illustrating the prominent role of glucose metabolism in NCC adaptation to physical changes in the environment. Thus, contrary to the general assertion that OXPHOS has a bioenergetic role in differentiated cells and that aerobic glycolysis has a biosynthetic role in proliferating non-differentiated stem cells (Ito and Suda, 2014; Shyh-Chang et al., 2013), NCC development relies on the concerted and sustained mobilization of all metabolic pathways dependent on glucose metabolism, i.e. glycolysis, OXPHOS, and PPP, cooperating together (Fig. 7). These findings illustrate the necessity for the NCC population to make the most of the potential of the carbon metabolism route, so that they are endowed with the capacity to meet the high metabolic demands suitable for the coordinated execution of diverse cellular events, in a minimum of time and in a rapidly evolving environment.

**Fig. 7.**
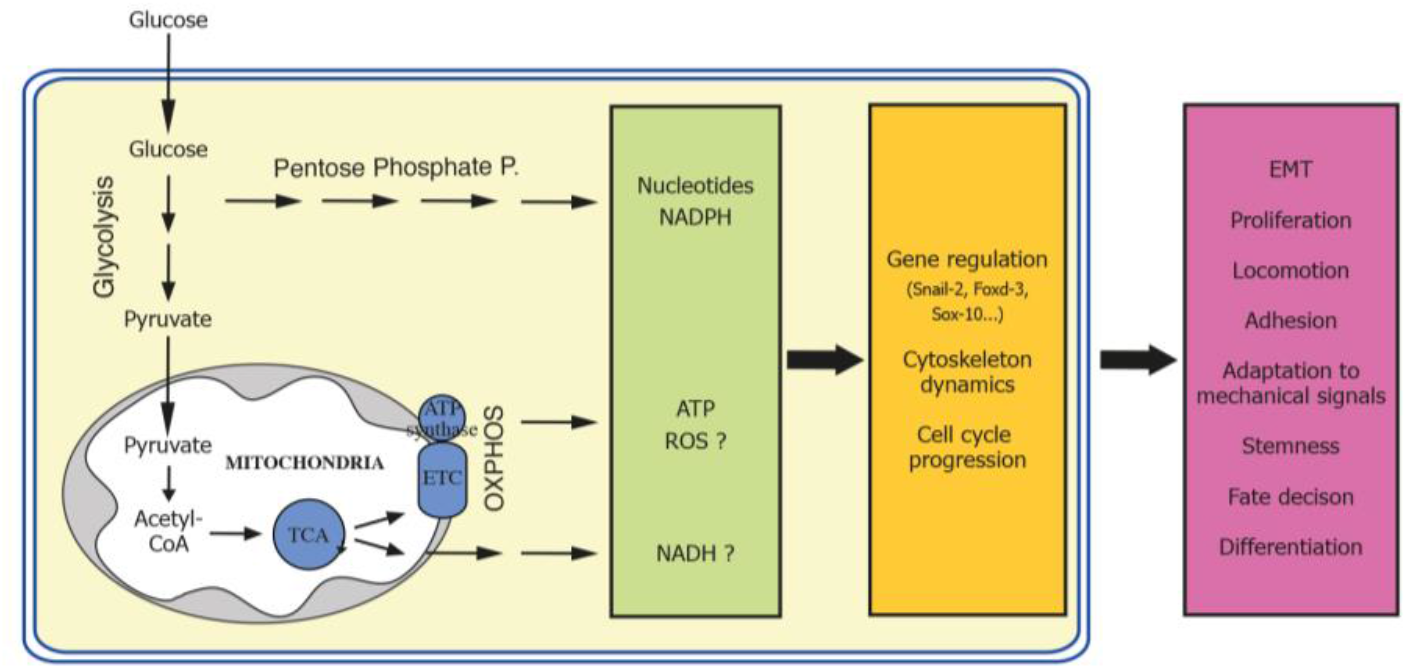
Schematic representation of the integration of carbon metabolism in NCC development. See discussion for details.

How NCC maintain balanced activities of the different metabolic pathways is currently unknown. This may be achieved through regulated levels of metabolic enzymes and products of each pathway, under the control of intrinsic genetic programs as well as extrinsic factors, including nutrient availability and demand (Lempradl et al., 2015). This view is supported by our observations as well as a recent report (Oginuma et al., 2017) that glycolytic enzymes are not uniformly expressed in the embryo at the time of NCC dispersion. Alternatively, the uncoupling protein UCP2, the ATPase inhibitory factor IF1, and the pyruvate dehydrogenate kinase PDK, known to suppress coupling between glycolysis and OXPHOS or to block mitochondrial activity, may contribute to metabolic rewiring in pluripotent stem cells (Sánchez-Aragó et al., 2013; Takubo et al., 2013; Zhang et al., 2011). It will be of interest to investigate whether expression of these players is subjected to spatiotemporal regulation in NCC.

We found no evidence for a significant contribution of a Warburg effect to trunk NCC development. Indeed, ECAR assessments and lactate measurements both revealed low lactate production in NCC, demonstrating that glucose is not used for ATP production through lactate production, but serves to fuel OXPHOS, PPP, and possibly amino acid pathways for biosynthetic purposes. Moreover, the fact that oligomycin did not increase lactate levels indicate that glucose metabolism cannot be rewired from OXPHOS to aerobic glycolysis in NCC. Our conclusion contrasts with a recent report proposing that, similarly to cancer cells, cranial NCC display a Warburg effect (Bhattacharya et al., 2020). The difference in metabolic activity between cranial and trunk NCC is likely to reside in the local O_2_ concentration in their environment. As for cancer cells in which hypoxia is known to promote metastatic spread (Semenza, 2012), cranial NCC reside in a hypoxic milieu, which favors prolonged NCC production from the NT through maintenance of EMT gene expression. Exposure to high O_2_ levels or knockdown of the hypoxia-inducible factor HIF-1 cause strong reduction of *Snail-2* expression and attenuation of NCC production (Scully et al., 2016). Trunk NCC dispersion in contrast is intimately associated with blood vasculature (Schwarz et al., 2009; Thiery et al., 1982), and it is not affected by high O_2_ levels or by deletion of the HIF-1 gene (Iyer et al., 1998; Morriss and New, 1979). Our data therefore indicate that trunk NCC display metabolic requirements distinct from their cranial counterparts and reveal that, although they possess in common with metastatic cells a number of cellular and molecular features, such as signaling pathways, EMT, migration properties and gene circuits, they do not share with them the same metabolic status. This reflects the strong adaptability of NCC to a great variety of environmental conditions throughout development, at the origin of the diversity in their migratory behaviors and fates.

We could not attribute to any metabolic pathway a specific role in the basic cellular processes, adhesion, locomotion and division. For example, despite PPP having a well-recognized role in nucleotide synthesis and maintenance of cellular redox balance (Patra and Hay, 2014), this pathway was not recruited solely for NCC proliferation but significantly contributed to cell velocity and persistence of movement as well. Likewise, OXPHOS and ATP production in mitochondria were not exclusively aimed at supporting active migration, but also played a key role in EMT and proliferation. A likely explanation is that these processes are extremely demanding in both bioenergetic and biosynthetic supplies (Zanotelli et al., 2021)(Salazar-Roa and Malumbres, 2017). Another reason is that enzymes, metabolites and byproducts of metabolic pathways are known to influence cellular programs independently of their canonical bioenergetics and biosynthetic roles (Miyazawa and Aulehla, 2018). NADH produced through TCA protects cells from oxidative stress (Bigarella et al., 2014; Khacho et al., 2016). This is illustrated by our observation that Rot-AA exhibit a stronger effect on NCC than oligomycin. Likewise, similarly to neurons (Herrero-Mendez et al., 2009), the use of the PPP for metabolizing glucose may ensure NADPH production for the maintenance of a cellular redox balance. TCA cycle intermediates, in particular acetyl-CoA and α-ketoglutarate, are involved in epigenetic regulation through histone acetylation and methylation (Kaelin and McKnight, 2013). Regulation of gene expression during NCC formation is under tight epigenetic control, involving DNA methylation, histone modifications, and ATP-dependent chromatin remodelers (Hu et al., 2014). Our finding of complete reprogramming of gene circuits driving entire developmental programs such as delamination and maintenance of stemness upon manipulating nutrient availability and metabolic pathways in NCC is therefore in favor of a direct role of cellular metabolism in regulation of nuclear transcription programs in NCC through epigenetic regulation.

An intriguing finding in our study is related to the effect of pyruvate on NCC differentiation and fate decision. Consistent with previous studies showing that mitochondrial pyruvate metabolism represses stem cell state and proliferation (Schell et al., 2017), we found that providing pyruvate to NCC in absence of glucose caused cell cycle arrest and loss of stem cell markers, but contrary to what has been reported for many other embryonic and stem cells (Ito and Suda, 2014; Khacho et al., 2016; Khacho and Slack, 2017; Shyh-Chang et al., 2013), it biased differentiation to a limited range of cell types where myofibroblasts predominate. In the embryo, myofibroblasts do not normally derive from trunk NCC but are only provided by cranial NCC. However, in *ex vivo* cultures in the presence of various hormones and growth factors (Dupin et al., 2018), myofibroblasts were recorded in trunk NCC progeny, together with skeletogenic and adipogenic cells. These data have been interpreted as a dormant capacity of trunk NCC to differentiate into non-neural derivatives normally obtained with cranial NCC. Our observation that myofibroblastic differentiation occurs essentially in pyruvate conditions in the complete absence of additives suggests that this dormant capacity can be uncovered by pyruvate, thereby illustrating the key role of metabolism in instructing NCC fate.

In conclusion, our data show that, contrary to pluripotent stem cells and cancer cells, NCC in the embryonic trunk region rely primarily on glucose oxidation through glycolysis and mitochondrial respiration for energy production and mobilize a large range of metabolic pathways downstream glucose uptake to meet their important bioenergetics and biosynthetic needs as well as to instruct their gene circuits driving their behavior and fate. These findings therefore reveal the intricate integration of cellular metabolism to basic cellular processes underlying cell behavior. How these metabolic pathways are regulated to accommodate the different steps of NCC formation both temporally and spatially remains to be investigated.

## MATERIALS AND METHODS

### Reagents

Bovine plasma fibronectin (1mg/ml stock) was from Sigma (cat. no. F1141). Dispase II at 5 U/ml stock solution in Hanks’ balanced saline was from Stemcell Technologies (cat. no. 07913). The following chemicals were prepared and used according to the manufacturers’ guidelines: 2-deoxy-D-glucose (2-DG, Sigma, cat. no. D8375), Na-iodoacetate (Sigma, cat. no. I2512), oligomycin-A (Sigma, cat. no. 75351), rotenone (Sigma, cat. no. R8875), antimycin-A (Sigma, cat. no. A8674), FCCP (Sigma, cat. no. C2920), 6-aminonicotinamide (6-AN, Cayman Chemicals, cat. no. 10009315), and UK-5099 (Tocris Biosciences, cat. no. 4186). 2-DG was prepared as a 1 M stock solution in glucose-, pyruvate- and glutamine-free DMEM and used at 5-10 mM; Na-iodoacetate was prepared as a 15 mM stock solution in H_2_O and used at 15 μM; oligomycin, rotenone, antimycin-A, and FCCP were prepared in DMSO as 100 μM stock solutions and used at 1 μM; 6-AN was prepared as 100 mM solution in DMSO and used at 100 μM, except in Seahorse analyses where it was used at 500 μM. UK-5099 was prepared as 100 mM solution in DMSO and used at 100 μM.

### Cell Culture

#### Generation of NCC primary cultures

NCC cultures were produced from NT explants obtained from quail embryos (Coturnix coturnix japonica) as described (Duband et al., 2020). Briefly, fertilized eggs purchased from a local farm (La Caille de Chanteloup, Corps Nuds, France) were incubated at 37-38°C for 54 h until the embryos reached stages corresponding to the chick stages 13-15 of Hamburger and Hamilton (HH) (1951), i.e. comprising 19-25 somite pairs. The eggshell was opened with curved dissecting scissors and the yolk transferred into phosphate-buffered saline (PBS). The embryo was cut off from the yolk and transferred into PBS in an elastomer-containing dish. An embryo portion of about 750-μm long was excised at the level of the last 5 somites with a scalpel under a stereomicroscope and subjected to mild enzymatic digestion by treatment with dispase II at 2.5 U/ml for 5-10 minutes at room temperature. The NT was dissected out manually using fine dissection pins under a stereomicroscope, freed from the surrounding tissues, and transferred for 30-60 minutes in DMEM medium (no glucose, no pyruvate, Gibco, cat. no. 11966025) supplemented with 0.5% fetal bovine serum for recovery from enzyme treatment. NT were explanted onto culture dishes or coverglasses previously coated with fibronectin at 10 μg/ml in PBS (i.e. about 5 μg/cm^2^) for a minimum of 1 h at 37°C. To ensure rapid initiation of NCC migration, NT were positioned with their dorsal side oriented down toward the substratum. Explants were cultured at 37°C under normoxic conditions in a humidified 5%-CO_2_ incubator in DMEM containing 1% serum, 100 U/ml penicillin, 100 μg/ml streptomycin, and 2 mM glutamine, and supplemented or not with 5 mM glucose and 1 mM pyruvate. The choices of the culture dish, culture medium, and duration of the culture were determined according to the purpose of the experiment and the method of analysis used (see below). Throughout each experiment, the morphology of the NT explant, area and progression of the NCC outgrowth, as well as individual cell shape and viability were evaluated, imaged, and assessed regularly under an inverted phase contrast Nikon microscope equipped with 6.3X, 10X, and 20X objectives.

#### Delamination assay

NT explants were cultured for 5 h, a time sufficient to allow NCC segregating from the dorsal NT to adhere to the substrate. NT was then removed manually from the dish using fine dissection pins under a stereomicroscope or by gentle flushing of culture medium with a pipet tip, to uncover delaminated cells.

### Cellular Bioenergetic Analyses

Bioenergetic profiles of NCC primary cultures were determined using a Seahorse Bioscience XF24 Analyzer. A single NT explant was deposited precisely at the center of each well of 24-well Seahorse plate previously coated with fibronectin in culture medium at 37°C in a humidified 5%-CO_2_ incubator, and NCC were allowed to undergo migration. Before preparation of the plate for Seahorse analysis, the areas of the NT explants were measured for normalization. Cultures were then rinsed in Seahorse XF medium (Agilent, cat. no 103575-100) without serum and incubated for 1 h at 37°C in normal atmosphere. Following incubation, the Seahorse assay was run according to the manufacturer’s instructions. Oxygen consumption rate (OCR) and extracellular acidification rate (ECAR) values, as readouts of basal mitochondrial respiration and glycolysis respectively, were assessed regularly every cycle (mix, wait and measurement during 3 minutes), the basal measurement consisting of 4 cycles and drug injections, 3 cycles during 30 minutes. The key parameters of mitochondrial respiration (ATP-linked respiration, maximal respiratory capacity and proton leak) were measured by means of a MitoStress test after sequential additions of oligomycin (1 μM), FCCP (0.75 μM), and Rot-AA (1 μM) through 3 cycles of measurements in 30 minutes. When appropriate, NCC primary cultures were processed after Seahorse analysis for immunolabeling, qPCR analysis, or intracellular ATP level and lactate measurement.

#### ATP and lactate measurements

Intracellular ATP and extracellular lactate levels were measured for each NCC primary cultures after Seahorse analysis using an ATPlite Bioluminescence assay kit from PerkinElmer (cat. no. 6016943) and the Lactate Fluorometric Assay kit from BioVision (cat. no. K607), respectively, according to the manufacturers’ instructions.

### Fluorescence Labeling of NCC Cultures

For immunolabeling, the following primary antibodies were used: Rabbit monoclonal antibody (mAb) to Snail-2 (clone C19G7, Cell Signaling, 1/300), mouse mAb to Sox-10 (Clone A2, Santa Cruz, 1/200), mouse mAb to β-catenin (clone 14, BD-Transduction Laboratories, 1/200), mouse mAb to paxillin (Clone 165, BD-Transduction Laboratories, 1/100), mouse mAb to α-SMA conjugated to Cy3 (Clone 1A4, Sigma, 1/300), mouse mAb to βIII-tubulin conjugated to Alexa488 (Clone Tuj-1, R&D Systems, 1/500), mouse mAb to HNK-1 described previously ((Tucker et al., 1984), undiluted culture supernatant), mouse mAb to MITF (clone C5, Abcam, 1/200), and mouse mAb to Glut-1 (Ab14683, Abcam, 1/50). Filamentous actin was detected using Texas Red™-X Phalloidin (Molecular probe, cat. no. T7471). NCC primary cultures were performed in 4-well plates or in 8-well Chambered Coverglass (Nunc, cat. no. 154511) coated with fibronectin, and fixed in 4% paraformaldehyde (PFA) in PBS for 15 minutes at room temperature for detection of all antigens, except for Snail-2 (5 minutes in 4% PFA), β-catenin (45 minutes in 1.5% PFA), and Glut-1 (see below). After permeabilization with 0.5% Triton X-100 in PBS for 15 minutes, cultures were blocked in PBS-3% BSA and subjected to immunofluorescence labeling using primary antibodies followed by incubation with appropriate secondary Ab conjugated to Alexa-fluor 488, Cy-3 or Cy-5 (Jackson Immunoresearch Laboratories), and processed for DAPI or Hoechst staining to visualize cells’ nuclei before mounting in ImmuMount medium (Shandon). For Glut-1 immunolabeling, unfixed NCC cultures were incubated with the Glut-1 antibody diluted in DMEM for 15 minutes at 15°C to avoid antibody internalization and immediately fixed in 100% ethanol at −20°C before rinsing in PBS and secondary antibody treatment. For labeling of mitochondria, MitoTracker Red (Invitrogen, cat. no. M7512) was added at 1 μM to unfixed cultures and incubated for 20-30 minutes at 37°C. After washes with 37°C pre-warmed PBS, cells were briefly fixed with 4% PFA and mounted. Preparations were observed with a Zeiss AxioImager M2 epifluorescence microscope equipped with 10X-63X fluorescence objectives (Acroplan 10X/0.25, 20X/0.45, Plan-Neofluar 40X/0.75 and 63X/1.25 oil) or with a Zeiss LSM 900 confocal microscope equipped with 10X-40X fluorescence objectives (Plan-Apochrome 40X/1.30 oil). Data were collected using the Zen or Airyscan2 software and processed using ImageJ software. For each series of experiments, images were acquired using equal exposure times and settings.

### In Situ Hybridizations for mRNA Detection on NCC Primary Cultures and Whole Mount Embryos

The following plasmids for mRNA probe synthesis were used: *Snail-2* from A. Nieto, *Foxd-3* from C. Erickson, *Sox-10* from P. Scotting, *Glut-1, PFK, PGK-1* and *PKM* from O. Pourquié. Linearized plasmid DNA was used to synthesize digoxigenin-UTP (Roche) labeled antisense probes with RNA polymerases from Promega and RNA probes were purified with Illustra ProbeQuant G-50 microcolumns (GE Healthcare). In situ hybridizations were performed either on NCC primary cultures produced in fibronectin-coated plastic dishes or on whole mount intact embryos collected at the appropriate developmental stages, using essentially the same procedure. Samples of embryos and cultures were fixed in 4% PFA in PBS for 2 h at room temperature or overnight at 4°C. They were hybridized overnight at 65°C with the digoxygenin-UTP-labeled RNA probes in 50% formamide, 10% dextran sulfate and Denhart’s buffer (0.5 μg probe/ml hybridization buffer) and washed twice in 50% formamide, 1x SSC and 0.1% Tween-20 at 65°C, then 4 times at room temperature in 100 mM maleic acid, 150 mM NaCl pH 7.5 and 0.1% Tween-20 (MABT buffer). After a 1-h pre-incubation in MABT buffer containing 10% blocking reagent (Roche) and 10% heat-inactivated lamb serum, samples were incubated overnight at room temperature with the anti-digoxygenin antibody (Roche). After extensive rinsing with MABT buffer, they were preincubated in 100 mM NaCl, 50 mM MgCl_2_, 1% Tween-20, and 25 mM Tris-HCl, pH 9.5, and stained with NBT-BCIP (Roche) following manufacturer’s guidelines. Preparations were observed and imaged with a Nikon stereomicroscope and data analyzed with ImageJ software.

### RNA Quantification by Quantitative RT-PCR Analyses

Total RNA of cells from cultures of 5-8 NT explants were extracted using the Ambion PureLink RNA Mini Kit (Invitrogen, cat. no. 12183018A) following the manufacturer’s guidelines. After quantification of RNA concentration, double-strand cDNA was synthesized from RNA using SuperScript IV reverse transcriptase (Invitrogen, cat. no. 18090050). Quantitative real time PCR were performed using the Power Syber Green Master Mix (Applied Biosystems, cat. no. 4368708) in a StepOne Plus RT-PCR apparatus (Applied Biosystems). Gene expression was assessed by the comparative CT (ΔΔCt) method with β-actin as the reference gene. The following primers were used: *Snail2fwd* GATGCGCTCGCAGTGATAGT; *Snail2rev* AGCTTTCATACAGGTATGGGGATA; *Sox10fwd* CGGAGCACTCTTCAGGTCAG; *Sox10rev* CCCTTCTCGCTTGGAGTCAG; *Foxd3fwd* TCTGCGAGTTCATCAGCAAC; *Foxd3rev* TTCACGAAGCAGTCGTTGAG; *Glut1fwd* AAGATGACAGCTCGCCTGATG; *Glut1rev* AGTCTTCAATCACCTTCTGCGG; *PFKfwd* TTGGAATTGTCAGCTGCCCG; *PFKrev* TGCAGACAACTTTCATAGGCATCAG; *βActinfwd* CTGTGCCCATCTATGAAGGCTA; *βActinrev* ATTTCTCTCTCGGCTGTGGTG.

### Cell Proliferation Assay

Cell proliferation in culture was monitored using the Plus EdU Cell proliferation kit for imaging from Life Technologies (cat. no. C10638). Briefly, NCC primary cultures were generated on fibronectin-coated coverglasses as for immunolabelings and were incubated with EdU at 20 μM in culture medium for 1 h at varying times of NCC development. Immediately after EdU incorporation, cultures were fixed in 4% PFA in PBS for 15 minutes at room temperature, permeabilized in 0.5 Triton-X100 for 15 minutes and treated for EdU detection using Alexa-555 Fluor azide in accordance with the manufacturer’s guidelines. After DNA staining with Hoechst, cultures were optionally processed for immunostaining and analyzed as described for immunolabelings.

### Cell Locomotion Assays

NCC primary cultures were performed in 8-well Chambered Coverglass coated with fibronectin. Up to 4 NT explants were distributed separately into each well, and were maintained at 37°C in a humidified 5%-CO_2_ incubator for about 2 h until the NT adhered firmly to the dish and NCC initiated migration on the substratum; then the cultures were transferred into a heated chamber (Ibidi) with a humid atmosphere containing 5% CO_2_/95% air placed on the motorized stage of a Leica DMIRE2 microscope equipped with a CoolSNAP HQ camera (Roper Scientific). Time-lapse video microscopy was performed with a 10x objective and phase contrast images were captured every 5 minutes during 16-24 h using the Micromanager software. Trajectories and positions of individual NCC in several explants recorded in parallel were tracked using Metamorph 7 software. The velocity of each cell was calculated as the ratio between the total length of its trajectory and the duration of the acquisition time and the persistence of movement as the ratio between the linear distance from the cell’s initial to final positions and the total distance covered by the cell. Evolution of speed and persistence every hour were determined. The progression of the migratory front of the NCC population every hour was measured using a custom GUI written in Matlab®(MathWorks®, Natick, Massachusetts, USA).

### Preparation of polyacrylamide gels

Polyacrylamide (PAA) gels were prepared on 32-mm glass coverslips and 8-well Chambered Coverglass for AFM and TFM experiments, respectively. The surface on which the PAA gels were polymerized was activated by immersion into a solution of 3-methacryloxypropyltrimethoxysilane (0.3%, Bind-Silane, Sigma, cat. no. 440159), 10% acetic acid aqueous solution (3%) in absolute ethanol during 3 minutes to enhance gel adhesion, washed 3 times with ethanol, and air-dried for 45 minutes. Thin sheets of PAA gel of different elastic properties were prepared from concentrations of acrylamide/bisacrylamide 5%/0.225% and 3%/0.08%, supplemented with 0.5% ammonium persulfate and 0.05% tetramethylethylenediamine (Sigma) in H20. Practically, for AFM experiments, 81 μl of these different mixtures were placed onto the surface of a 32 mm-diameter coverslip and covered with 14 mm-diameter coverslip. For immunostaining and video microscopy experiments, 2.5 μl of the mixtures were placed in the wells of 8-well chambered Coverglass and covered with a 6 mm-diameter coverslip. Top coverslips were treated previously with Repel-Silane ES (Merck, cat. no. GE17-1332-01) during 5 minutes to prevent PAA gel adhesion. After 30 minutes of polymerization, the top coverslips were removed and the PAA gels were rinsed 3 times with PBS, treated twice for 5 minutes with sulfo-SANPAH, UV-photoactivated with Bio-Link Crosslinker BLX-E254 (Biotech), and rinsed 3 times with 50mM HEPES at pH 8.5. For traction force microscopy experiments, the gel mixtures contain FluoSpheres from Sigma (TM carboxylate modified, 0.2μm, red (580/605), cat. no. F8810), at 1/50 dilution. A solution of fibronectin was layered onto the PAA gels at 30μg/cm^2^ and incubated overnight at 4°C under agitation.

### AFM assessments on polyacrylamide gels and NCC

The elastic modulus for both PAA gels and NCC cultured for 4-6 h was assessed by AFM JPK NanoWizard Sence+ (Brucker, Billerica, Massachusetts, U.S.A) coupled to a Zeiss Axio-Observer z1 inverted microscope, as described previously (Ben Bouali et al., 2020) using μMash CSC38/NO AL probes (MikroMasch®, Sofia, Bulgaria). PAA gels and NCC were set in culture medium containing 25mM HEPES and maintained at 37°C. For PAA gels, measurements were achieved considering a measuring point grid with 0.4-μm meshes over a 4.8×4.8 μm^2^ surface set in gel center. For NCC, measurements were achieved considering a measuring point grid with 1-μm mesh over a 12×12 μm2 surface set on cells using visual inspection with a phase contrast microscope. Setting Poisson’s ratio ν = 0.5 for both PAA gels and NCC, AFM spectroscopy curves were analyzed according to (Bilodeau, 1992), for quadrilateral pyramid probe using a home written program in Matlab®. The elastic modulus of the PAA gels prepared with proportion of acrylamide/bisacrylamide 5%/0.225% and 3%/0.08% were of 6320 +/− 25 Pa and 1250 +/− 32 Pa (S.E.M), respectively, and were referred in the text as stiff and soft gels.

### TFM assessments

NT explants were cultured onto bead-embedded PAA gels in various nutrients for 4-6 h and transferred into a heated chamber with a humid atmosphere containing 5% CO_2_/95% air placed on the motorized stage of a Zeiss LSM 900 confocal microscope. A specific injector connected to 8-well Chambered Coverglass was designed for trypsin delivery and installed on the Chambered coverglass. Images of bead positions were acquired using Plan-Apochrome 40X/1.30 oil objective as follows: Images with force were acquired with 2-min interval leading to a 12-min observation time (stress images) and one image was acquired without force (reference image) taken after NCC have detached by trypsinization and no longer exerting forces on the substrate. Confocal stress and reference images were aligned using a Matlab®script. The gel deformation was measured by the PIV (Particle image velocimetry) technique using the ImageJ plug-in PIV to get the bead displacement field. Finally, the ImageJ plug-in FTTC (Fourier Transform traction cytometry) was used to measure the traction force, exerted by NCC and responsible of PAA gel deformation and bead displacement (Martiel et al., 2015). The number of NCC in the traction force field was measured by counting NCC nucleus stained with NucSpot®Live 650 and a confocal image was generated before starting TFM measurements. The stress taken into account is the mean stresses calculated from 6 couples of reference-stress image corresponding to 2, 4, 6, 8, 10 and 12 minutes TFM assessments. It was observed that between 2 and 12 minutes calculated stress was almost constant. The stress per cell is the stress divided by the number of nuclei. The force per cell was calculated by multiplying the stress/cell by the area of the interrogation window used in the PIV.

### Energy estimation

We estimated produced energy during NCC displacement on soft and stiff gels by calculating the product of the average force F obtained by TFM and the average displacement D per hour obtained by video microscopy tracking during the time frame of TFM studies. S.D. of the product F by D was calculated based on the following formula (Goodman, 1960): σ (F×D) = √ (〖E(F)〗 ^2 〖σ(D)〗 ^2+ 〖E(D)〗 ^2 〖σ(F)〗 ^2+ 〖σ(F)〗 ^2 〖σ(D)〗 ^2), with σ the S.D. and E(F) and E(D) mean value of F and D respectively, assuming F and D with normal distribution. To evaluate the statistical significance of two energy values, i.e., *m*_1_ ± *σ*_1_ and *m*_2_ ± *σ*_2_, we calculated Student’s *t* and degrees of freedom, *n*_1_ + *n*_2_ – 2, assuming *n*_1_ = *n*_2_ = 5 for size of and *m*_2_, to obtain p values. For this, we considered 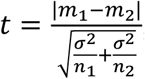, and *σ* common S.D, given by 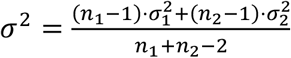.

### Statistical Methods

Statistical analyses were performed using Prism 7 (GraphPad). Unless specified, multiple comparisons were done respective to the control in the same experimental conditions. For statistical analysis of data we used One-way ANOVA parametric test after the validation of normality and equality of variances using Shapiro-Wilk and Brown-Forsythe methods, respectively. Otherwise, non-parametric Krustal-Wallis test was used. For comparison between two conditions we use unpaired t-test two-tailed test after the validation of normality and equality of using Shapiro-Wilk and equality of variances. Otherwise, non-parametric Mann-Whitney test was used. For statistical analysis of qPCR results, the data obtained for each drug or nutrient condition were compared to that of the control medium using unpaired t-test two-tailed test. Unless specified, at least three independent experiments were carried out for each procedure. Each NT explants was considered as an individual sample. The n of samples analyzed is indicated in each graph. Data are expressed as mean values ± S.D. or S.E.M and the P values are * p < 0.05, ** p < 0.01; *** p < 0.001; **** p < 0.0001. Results are considered statistically significantly different when p < 0.05.

## ACKNOWLEDGMENTS

We deeply thank Chantal Thibert and Sakina Torch for providing advice in cellular metabolism and critical reading of the manuscript and Frédéric Relaix for constant support. Special thanks to Olivier Pourquié and Masayuki Oginuma for providing cDNA probes for chicken glycolytic enzymes. We thank Xavier Decrouy and the IMRB imaging platform.

## AUTHOR CONTRIBUTIONS

Nioosha Nekooie-Marnany performed experiments, analyzed data and contributed to the manuscript. Redouane Fodil provided expertise in biomechanics and performed experiments for AFM and TFM. Sophie Féréol provided expertise in biomechanics and prepared the PAA gels. Marine Depp performed NCC primary culture experiments. Roberta Foresti and Roberto Motterlini provided expertise in cellular metabolism and Seahorse technology and revised the manuscript. Jean-Loup Duband conceived and designed the experimental approach, performed experiments, analyzed the data, and wrote the manuscript. Sylvie Dufour conceived and designed the experimental approach, performed experiments, analyzed the data, and contributed to the manuscript.

